# Glutaminolysis promotes the function of follicular helper T cells in lupus-prone mice

**DOI:** 10.1101/2024.11.25.625088

**Authors:** Seung-Chul Choi, Yong Ge, Milind V. Joshi, Damian Jimenez, Lauren T. Padilla, Cassandra LaPlante, Jeffery C. Rathmell, Mansour Mohamadzadeh, Laurence Morel

## Abstract

Glutamine metabolism is essential for T cell activation and functions. The inhibition of glutaminolysis impairs Th17 cell differentiation and alters Th1 cell functions. There is evidence for an active glutaminolysis in the immune cells of lupus patients. Treatment of lupus-prone mice with glutaminolysis inhibitors ameliorated disease in association with a reduced frequency of Th17 cells. This study was performed to determine the role of glutaminolysis in murine Tfh cells, a critical subset of helper CD4^+^ T cells in lupus that provide help to autoreactive B cells to produce autoantibodies. We showed that lupus Tfh present a high level of glutamine metabolism. The pharmacological inhibition of glutaminolysis with DON had little effect on the Tfh cells of healthy mice, but it reduced the expression of the critical costimulatory molecule ICOS on lupus Tfh cells, in association with a reduction of autoantibody production, germinal center B cell dynamics, as well as a reduction of the frequency of atypical age-related B cells and plasma cells. Accordingly, profound transcriptomic and metabolic changes, including an inhibition of glycolysis, were induced in lupus Tfh cells by DON, while healthy Tfh cells showed little changes. The T cell-specific inhibition of glutaminolysis by deletion of the gene encoding for the glutaminase enzyme GLS1 largely phenocopied the effects of DON on Tfh cells and B cells in an autoimmune genetic background with little effect in a congenic control background. These results were confirmed in an induced model of lupus. Finally, we showed that T cell-specific *Gls1* deletion impaired T-dependent humoral responses in autoimmune mice as well as their Tfh response to a viral infection. Overall, these results demonstrated a greater intrinsic requirement of lupus Tfh cells for their helper functions, and they suggest that targeting glutaminolysis may be beneficial to treat lupus.

## Introduction

The non-essential amino acid glutamine is required for T cell proliferation, serving as a precursor for nucleotide synthesis. In the mitochondria, glutamine is degraded by a process called glutaminolysis in which glutamine is converted to glutamate by glutaminases, including glutaminase 1 encoded by *Gls1*, then into alpha-ketoglutarate (αKG), which enters the TCA cycle and contributes to oxidative phosphorylation (OXPHOS) and ATP production. Glutamate is also a precursor for the biosynthesis of glutathione, which is essential for redox homeostasis as well as non-essential amino acids. αKG is a critical regulator of chromatin accessibility by serving as a co-factor of histone and DNA demethylases^1^.

Glutaminolysis has been investigated in T cells largely through targeting *Gls1*, either pharmacologically or genetically. *Gls1* deletion in T cells impairs Th17 cell differentiation and alters Th1 cell functions, in association with epigenetic changes^2^. ICER, a transcription factor that promotes Th17 differentiation, directly transactivates *Gls1* expression^3^, thus inducing the metabolic program that supports Th17 cells. PBMCs from patients with systemic lupus erythematosus (SLE) and splenocytes from the lupus-prone MRL/Lpr mice present high levels of glutamate and *GLS1* expression^4^. Treatment of MRL/Lpr mice with the *Gls1* inhibitors CB839^4^ or BPTES^5^ ameliorated disease by reducing the frequency of Th17 cells, a major pathogenic subset in this model. Paradoxically, *Gls1* inhibition reduced glycolysis, which is required for Th17 differentiation^6^, by decreasing the expression of HIF1α^5^. Other mechanisms by which glutaminolysis contributes Th17 differentiation have been identified in other models^2^. Elevated expression of *Gls1* in psoriasis promotes the expression of IL-17a by enhancing histone acetylation at the *Il17a* promoter^7^. In addition, the glutamine flux through the TCA cycle and the accumulation of glutaminolysis intermediates that influence chromatin accessibility are tightly controlled in Th17 cells by LBK1 that senses mitochondrial integrity^8^. Glutamine-driven OXPHOS also increases apoptosis resistance and cytotoxicity of Th17 cells against tumors, which could otherwise not be achieved by glycolysis^9^. Pharmacological inhibition of glutaminolysis with 6- diazo-5-oxo-L-norleucine (DON) prevented allograft rejection by reducing T cell activation^10^. On the other hand, DON treatment upregulated oxidative phosphorylation (OXPHOS) in tumor infiltrating CD4^+^ T cells and promoted an effector long-lived phenotype^11^.

Glutaminolysis contributes critically to functional B cell differentiation, with differentiation of human B cells into plasmablasts supported by TLR9 signaling and type I interferon (IFN) to orchestrate glutamine-dependent OXPHOS^12^. Similar to CD4^+^ T cells^13^, B cells from SLE patients are characterized by an enhanced OXPHOS^14^. Since activation through TLR9 ligands and type I IFN is a characteristic feature of lupus B cells, these results suggest that targeting glutaminolysis may reduce plasmablast differentiation and autoantibody production in SLE. However, treatment of MRL/lpr mice with CB839 did not affect their plasma cells^4^. Glutaminolysis is also required for the production of IL-10 by human B10 cells and their suppressive activity^15^, but the frequency of B10 cells increased in MRL/Lpr mice treated with CB839^4^. Thus, the role of glutaminolysis in B cells is complex and may be context-dependent.

We recently investigated the role of glutaminolysis in the expansion of follicular helper CD4^+^ T (Tfh) cells and germinal center (GC) B cells in lupus and the resulting autoantibodies in the lupus-prone triple congenic (TC) B6.*Sle1*.*Sle2*.*Sle3* mouse model. We have shown that treatment of TC mice with DON strongly reduced anti-dsDNA IgG, virtually eliminated GCs and reduced the frequency of GC B cells^16^. *In vitro*, however, DON did not alter the differentiation of purified TC B cells into antibody-producing cells under either T-dependent and T-independent conditions,^17^ suggesting that DON may exert an extrinsic effect on B cells *in vivo*. DON treatment of TC mice, did not reduce the frequency of Tfh cells, but reduced their ICOS and BCL-6 expression^16^. In addition, DON did not reduce the Tfh polarization of either TC or healthy control purified CD4^+^ T cells *in vitro*^18^. A slight reduction in the frequency of Tfh cells and GC B cells was observed in MRL/lpr mice treated with CB839^4^. These results suggest that glutaminolysis is not required for Tfh cell differentiation, but it may affect their function, as it alters Th1 function^2^.. In support of this hypothesis, we showed that TC Tfh cells exhibit a differential expression of amino acid transporters as well as an enhanced glutamate metabolism compared to healthy C57BL/6 (B6) controls^19^.

In this study, we further investigated the role of glutaminolysis in the expansion of Tfh cells and germinal center (GC) B cells in lupus mice to further gain mechanistic insights. We analyzed the effect of the pharmacologic inhibition of glutaminolysis with DON on the transcriptome and metabolome of Tfh cells. We used the (NZW x BXSB.Yaa)F1 mouse model of the disease, hereafter referred to as W.Yaa, in which the *Yaa* duplication of the *Tlr7* gene controls pathogenesis largely through TLR7/type 1 IFN signaling^20^, a signaling pathway critically involved in SLE^21^, compared to healthy control B6 mice. TLR7 signaling in W.Yaa mice promotes an extrafollicular differentiation of autoantibody-producing cells, with a robust expansion of age-related B cells (ABCs) and extrafollicular T helper (Texfh) cells^22^, two subsets corresponding to the DN2 B cells^23^ and Tph cells^24^ expanded in SLE patients. The W.Yaa model is thus highly relevant to elucidate the mechanisms in rewiring cellular metabolism to potentially control autoreactive Tfh cells and thus mitigate disease progression. We have demonstrated that the pharmacological inhibition of glycolysis with 2-deoxyglucose (2DG) completely reversed the autoimmune manifestations and renal pathology in W.Yaa mice^25^. Here, we show that DON had little effect on B6 mice, but reduced the frequency of ABCs and plasma cells in W.Yaa mice. DON had no effect on the frequency of W.Yaa Tfh and Texfh cells, but significantly reduced the expression of ICOS and OX40, which are critical regulators of Tfh cell differentiation, maintenance and function^26,27^. DON also uniquely modulated the transcriptome and metabolome of W.Yaa Tfh cells. We then investigated the T cell-intrinsic effect of *Gls1* deficiency in T cells on Tfh and GC B cell responses in B6.*Sle1b* mice, a congenic strain derived from the NZM2410 lupus model carrying mutations in 6 SLAM family genes^28^, which functionally regulate Tfh cells and GC B cells^29, 30^. Consequently, B6.*Sle1b* mice produce anti-dsDNA IgG autoantibodies^31^ and robustly expend GC B cells^32^ and Tfh cells^33^. Here, we showed that the genetic inhibition of glutaminolysis in T cells reduced both spontaneous and induced autoantibody production. Furthermore, it uniquely impaired the response to protein immunization as well as to a viral infection in *Sle1b* mice. Thus, these results suggest that lupus CD4^+^ T cells have a higher intrinsic demand for glutaminolysis, which supports Tfh functions and indirectly promotes autoantibody production.

### DON reduced autoantibody production and selectively affected CD4^+^ T cell phenotypes in W.Yaa mice

Treatment of W.Yaa mice with DON reduced the production of anti-dsDNA IgG while it increased in untreated mice (Fig. 1A). Only a few W.Yaa mice presented a high frequency of GC B cells, and DON did not have a significant effect on their frequency (Fig. 1B). However, the higher dark zone to light zone (DZ / LZ) ratio of the GC B cells present in W.Yaa mice was reduced to B6 levels by DON (Fig. 1C, D), suggesting that DON altered the GC B cell dynamics. As expected with the *Tlr7* duplication, W.Yaa mice presented a large population of (ABCs, and their frequency was reduced by DON (Fig. 1E). Finally, DON reduced the frequency of splenic plasma cells (Fig. 1F). This could be a direct result of the reduced ABC population, as well as the altered GC B cell development.

**Figure 1.**
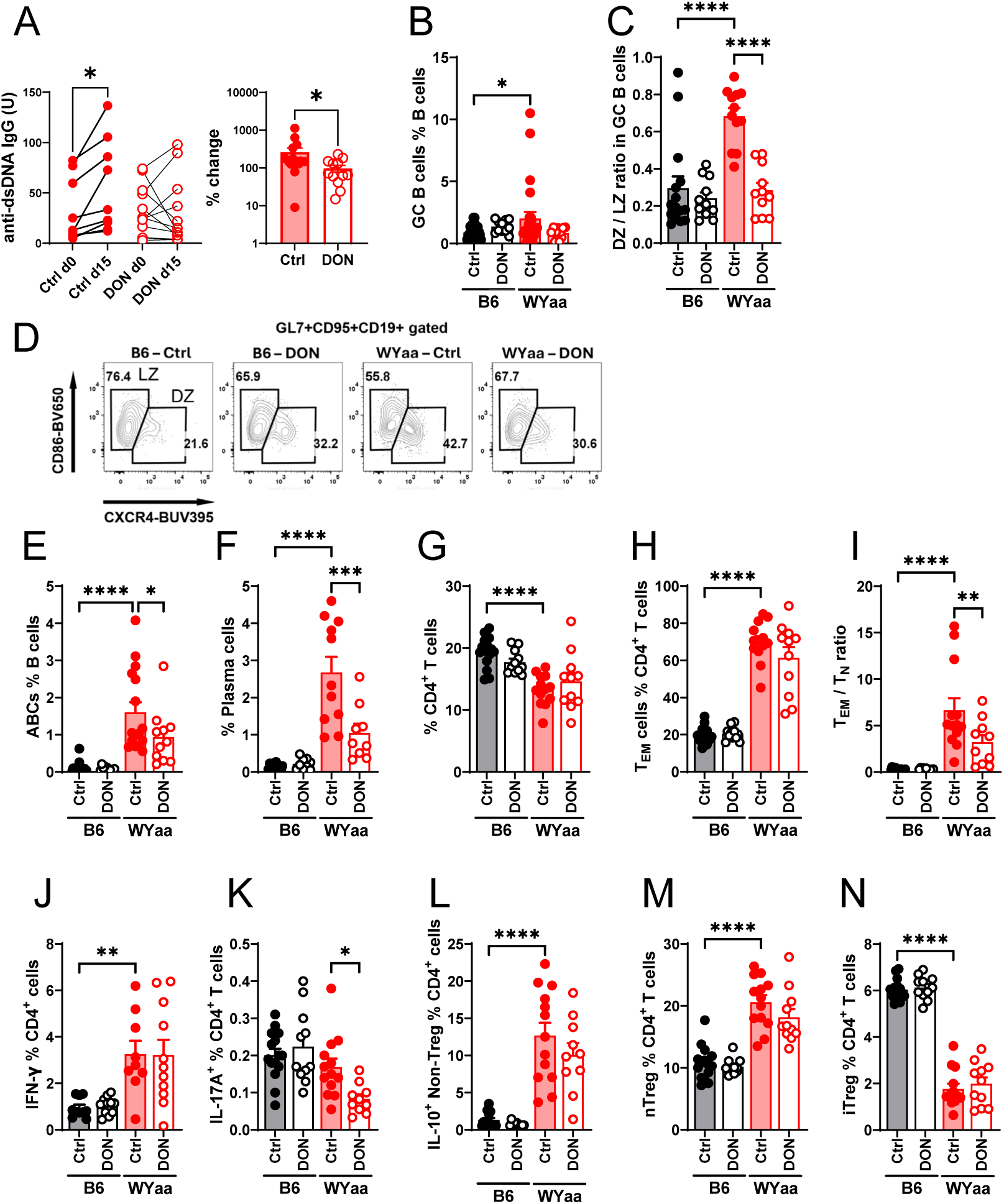
**DON reduced autoantibody production and selective CD4^+^ T cell activation**. **A**. Serum anti-dsDNA IgG measured before and at the end of treatment. Left: paired Unit values for each mouse, N = 9, paired t test. Right: Percent change after 2 weeks, N = 12 – 15. Means ± SEM, Mann Whitney test. **B**. Frequency of GC B cells **C**. GC B cell DZ/LZ ratio with representative FACS plots (**D**). Frequency of ABCs (**E**), plasma cells (**F**), total CD4^+^ T cells (**G**) and Tem cells (**H**). **I**. Tem/Tn ratio. Frequency of IFNγ^+^ (**J**), IL-17A^+^ (**K**), and IL-10^+^ (**L**) in FOXP3-negative CD4^+^ T cells. Frequency of nTreg (**M**) and iTreg (**N**) cells. Means ± SEM, N = 10 – 15 compared with Šídák’s multiple comparisons tests. *: P < 0.05; **: P < 0.01; ***: P < 0.001; ****: P < 0.0001.

W.Yaa mice present a decreased frequency of CD4^+^ T cells that did not changed with DON (Fig. 1G). There was a trend for DON reducing the high frequency of effector memory T cells (Tem) in W.Yaa mice leading to a significantly decreased Tem / naïve T (Tn) cell ratio, which is typically elevated in lupus-prone mice (Fig. 1H, I). DON had no effect on the high frequency of IFNγ- producing cells, but it reduced the small population of IL-17A-producing cells (Fig. 1J, K), which is consistent with the effect of glutaminolysis inhibition reported by others on Th1 and Th17 cells^2,5^. IL-10-producing FOXP3-negative (non-Treg) CD4^+^ T cells are expanded in lupus, including in W.Yaa mice, and the frequency of these cells was not affected by DON (Fig. 1L). Finally, W.Yaa mice present skewed frequencies of thymic-derived “natural” nTreg and peripherally induced iTreg cells, which were also not affected by DON (Fig. 1M, N).

As the TC lupus-prone model^16^, W.Yaa mice present an expanded population of Tfh cells as well as follicular regulatory T (Tfr) cells, with a skewed Tfh/Tfr ratio that were not altered by DON (Fig. 2A-C). However, as in TC mice, DON reduced the high expression of ICOS on Tfh and Tfr cells (Fig. 2D, E). Consistent with the high frequency of ABCs, W.Yaa mice develop a high frequency of extrafollicular helper CD4^+^ T cells (Texfh) and their corresponding FOXP3^+^ regulatory subset of Texfr cells, with a frequency not affected by DON (Fig. 2F, G). The Texfh / Texfr ratio was, however, reduced by DON (Fig. 2H), which could contribute to the lower ABC frequency (Fig. 1E). DON also reduced ICOS expression on Texfh and Texfr cells (Fig. 2I, J), as well as the expression of OX-40 on Tfh and Texfh cells (Fig. 2K, L). ICOS is an inducible costimulatory receptor that was downregulated by DON not only in Tfh-related subsets, but also in Tn, Tem and Treg cells (Fig. S1A-D), suggesting a requirement for glutaminolysis for ICOS expression. The downregulation of OX40 by DON was more selective to Tfh-related subsets, with no change observed Tn, Tem, and Treg cells (Fig. S1E-H), but a reduction in Tfr and Texfr cells (Fig. s1I, J) mirroring that of Tfh and Texfh cells (Fig. 2K, L). Contrary TC Tfh cells^16^, the expression of BCL-6 was not affected by DON (Fig. 2M). It should be noted that DON had no effect on B6 Tfh cells and related subsets for the parameters evaluated in this study.

**Fig. 2.**
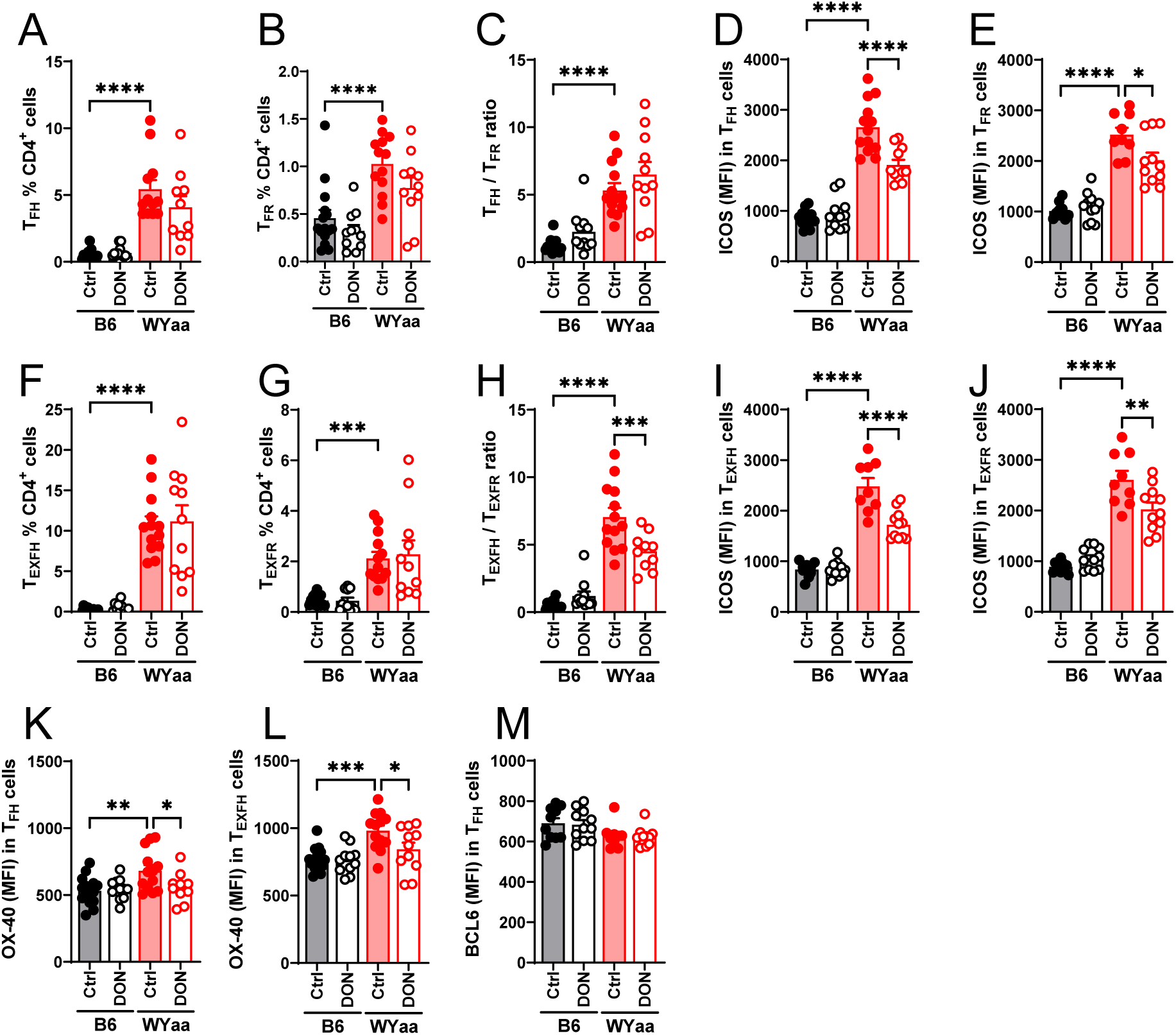
DON reduced the expression of ICOS and OX-40 on W.Yaa Tfh and Texfh cells with minimal effect on B6 cells. Frequency of Tfh (**A**) and Tfr (**B**) cells. **C.** Tfh/Tfh ratio. ICOS expression of Tfh (**D**) and Texfh (**E**) cells. Frequency of Texfh (**F**) and Tfr (**G**) cells. **H.** Tfh/Tfh ratio. ICOS expression of Tfh (**I**) and Texfh (**J**) cells. OX-40 expression on Tfh (**K**) and Texfh (**L**) cells. **M**. BCL-6 expression on Tfh cells. Means ± SEM, N = 10 – 15 compared with Šídák’s multiple comparisons tests. *: P < 0.05; **: P < 0.01; ***: P < 0.001; ****: P < 0.0001.

Overall, the DON treatment selectively altered W.Yaa CD4^+^ T cells, including the expression of critical co-stimulatory receptors on Tfh-related subsets, suggesting that glutaminolysis altered their maintenance and/or function, while it showed minimal effects on healthy B6 CD4^+^ T cells.

### Transcriptomic reprograming of W.Yaa Tfh cells by DON

To unbiasedly investigate the cellular responses to DON treatment, RNA sequencing analyses were performed on splenic Tfh cells isolated from DON-treated and untreated W.Yaa mice compared to DON-treated and untreated B6 mice. DON treatment resulted in a remarkable transcriptomic reprogramming of splenic Tfh cells isolated from both mouse strains, as demonstrate by principal component (PC) analysis (Fig. 3A). Accordingly, differentially expressed genes (DEGs) were identified in the DON-treated group and the control group (Fig. 3B). The main genes downregulated in W.Yaa Tfh cells included a number of heat shock proteins (e.g., *Hspa8, Hsph1, Dnaja1, Dnajb1*), which are generally triggered by cellular stress and inflammation. However, the expression of some of these genes, including *Hspa1a* and *Hspa1b*, were also activated in splenic Tfh cells derived from B6 mice after DON treatment. Furthermore, few DEGs were shared between W.Yaa and B6 Tfh cells (Fig. 3C), highlighting a specific effect of DON on autoimmune Tfh cells. Gene set enrichment analysis (GSEA) showed that DON mainly inhibited the signaling pathways related to the cell cycle and type I IFN response, particularly in W.Yaa Tfh cells (Fig. 3D). Here, the expression of a group of cell cyclins and cell division proteins as well as IFN-stimulated genes (ISGs), which demonstrate the highest expression in untreated W.Yaa Tfh cells, was reduced in W.Yaa Tfh cells by DON to the level of B6 Tfh cells (Fig. 3E). In addition, genes involved in glycolysis, including hexokinase 2 (*Hk2*), glucose transporter (*Slc2a1/Glut1*), and pyruvate kinase (*Pkm*), were also inhibited by DON. Lactate dehydrogenase A (*Ldha*), which critically supports glycolysis and T cell differentiation^34^, was also suppressed by DON, while its isoenzyme *Ldhb*, whose deletion in T cells enhances glycolysis, was increased (Fig. 3E), highlighting the optimization of glycolysis activity by DON to potentially control T cell differentiation.

**Fig. 3.**
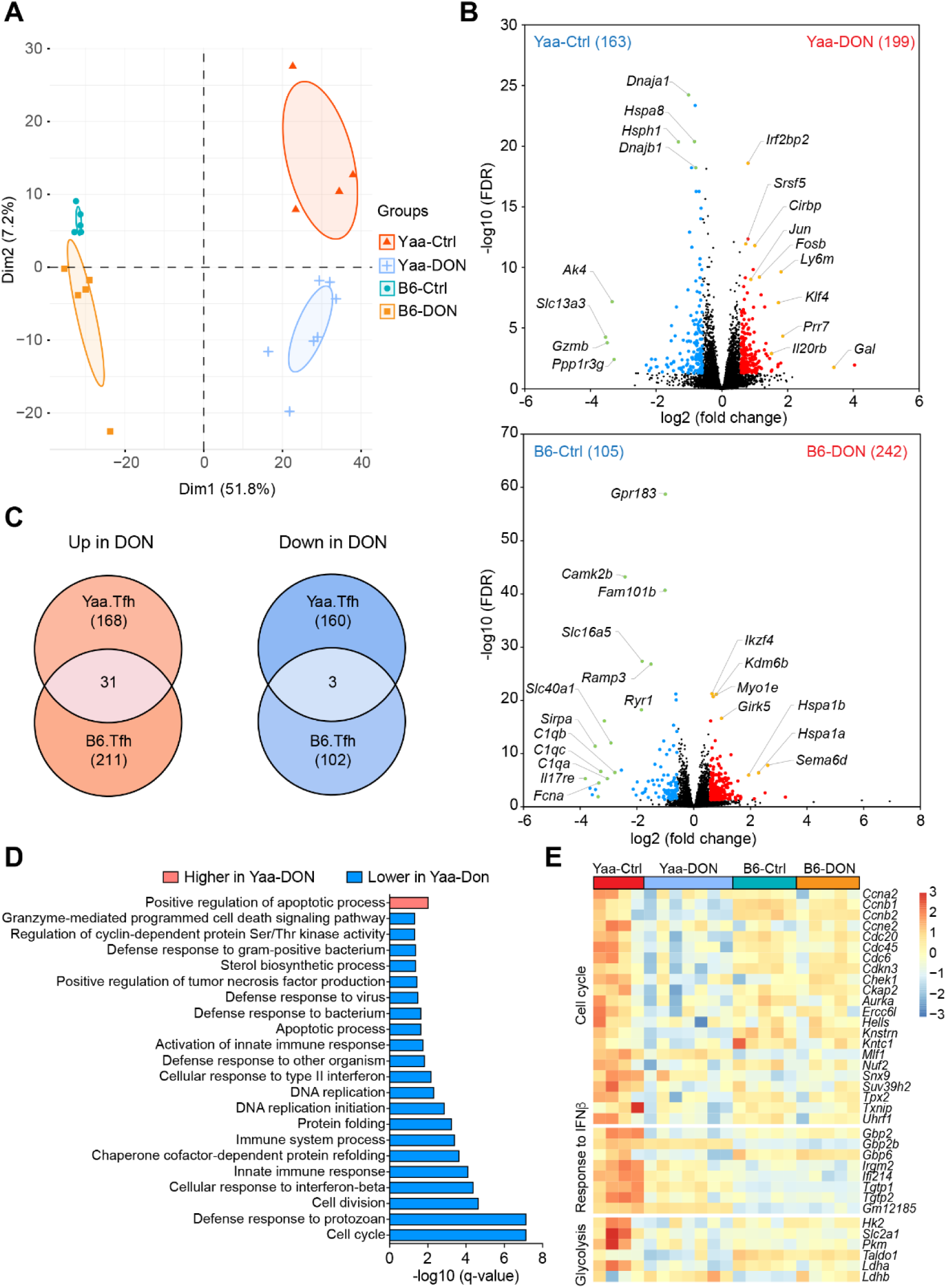
DON uniquely regulates the transcriptomic programming of W.Yaa Tfh cells. **A.** Principal component analysis (PCA) plot of the transcriptomes of Tfh cells isolated from W.Yaa and B6 mice treated with DON or untreated controls (n = 4-7/group). **B.** RNA-seq results depicted as volcano plots showing the differentially expressed genes (DEGs) (fold change > 1.5, FDR < 0.05) between DON and control groups. The number of DEGs upregulated in indicated groups is shown in parentheses. **C.** Venn diagrams depicting the unique impact of DON on W.Yaa Tfh cells. **D.** DAVID Gene Ontology (GO) analysis, related to biological process, showing upregulated or downregulated pathways in W.Yaa Tfh cells by DON treatment. **E.** Heatmap of DEGs related to cell cycle, response to IFNb, and glycolysis.

### DON uniquely altered the metabolism of W.Yaa Tfh cells

Next, we analyzed the metabolomes of splenic Tfh cells isolated from the same cohorts of W.Yaa and B6 mice as described above. While no segregation was seen in B6 Tfh cells, DON treatment significantly shifted the metabolome of W.Yaa Tfh cells toward a B6-like metabolic phenotype (Fig. 4A). In particular, amino acid (e.g., glutamate and arginine) and nucleotide metabolism was the primary metabolic pathway affected by DON (Fig. 4B). As expected, increased glutamine and decreased glutamate levels were evident in Tfh cells from DON-treated W.Yaa mice, but not in these cells from B6 mice (Fig. 4C and data not shown). g-Aminobutyric acid (GABA), which inhibits T cell autoimmunity and the development of inflammatory T cells^35^, was enriched by DON (Fig. 4C). One of the major functions of proliferating cells is to provide electron acceptors for aspartate biosynthesis^36^. Interestingly, aspartate as well as a number of nucleobases (cytosine, guanine and uracil), nucleosides (guanosine, inosine, uridine and cytosine) and nucleotides (uridine 5’-monophosphate and guanidine 5’-monophosphate) were strongly reduced in Tfh cells from DON-treated W.Yaa mice (Fig. 4C), potentially resulting in a global suppression of cellular metabolic activities as demonstrated by massive reductions in amino acids, including alanine, valine, proline, leucine, isoleucine, tyrosine, phenylalanine and tryptophan (Fig. 4C).

**Fig. 4.**
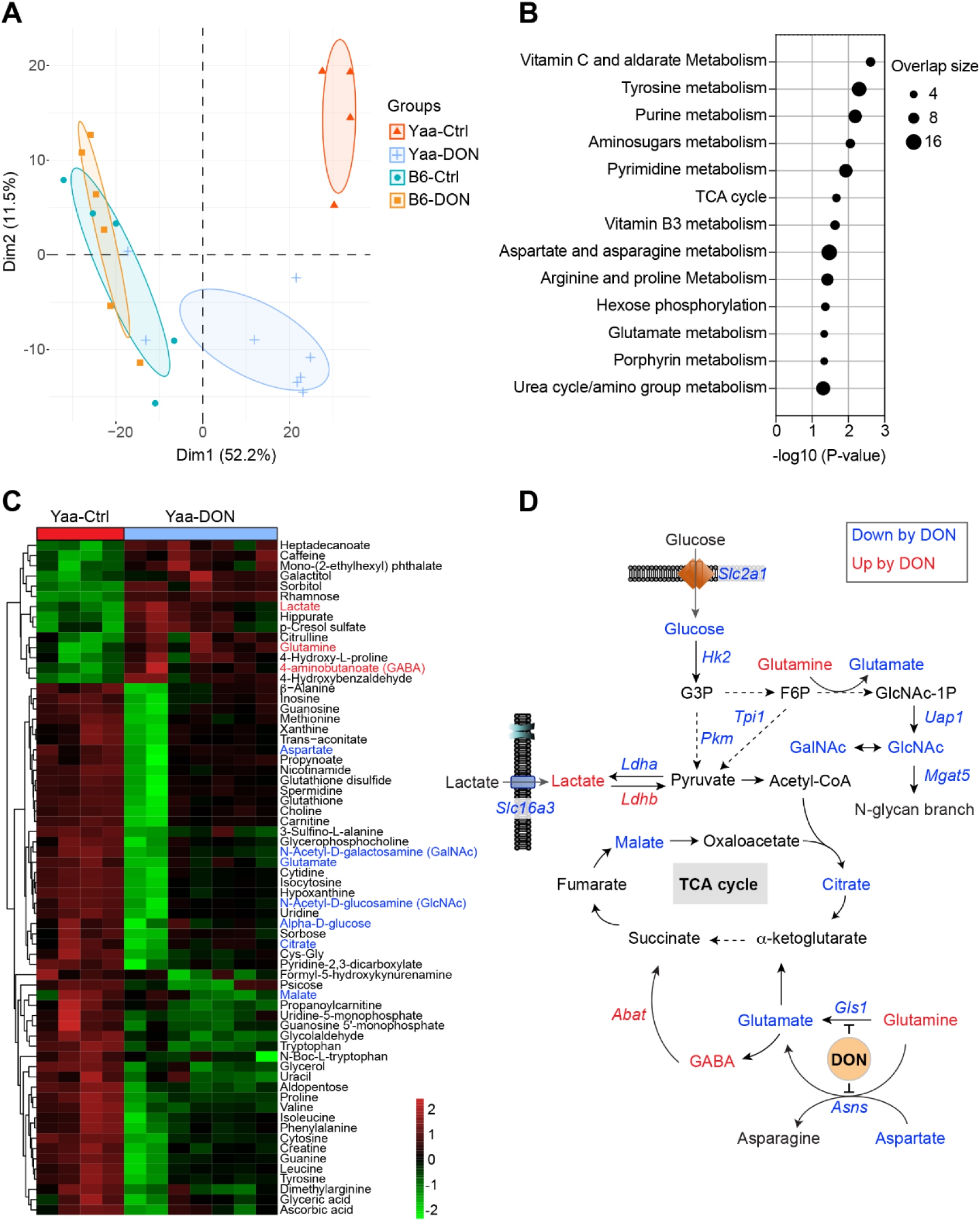
DON inhibits glycolysis and metabolically controls cell proliferation of autoimmune Tfh cells. **A**. PCA plot of metabolite features in Tfh cells isolated from W.Yaa and B6 mice after DON treatment versus untreated controls (n=4-7/group). **B**. Significant metabolic pathways in Tfh cells from W.Yaa mice treated with DON and control. Note: no significant pathway was identified between DON-treated and untreated B6 mice. **C**. Heat map of differentially enriched metabolites in W.Yaa Tfh cells. **D.** Transcriptional and metabolic integration illustrating the suppression of glycolysis and *N*-glycosylation activities in W.Yaa Tfh cells by DON. Gene (italic) and metabolites with higher and lower levels in DON group compared to controls are highlighted in red and blue, respectively, and shown on the heatmap in (C).

Taken together, transcriptional and metabolic integration uncovered a marked inhibition of glycolysis in autoimmune Tfh cells by DON, which was accompanied by a slight reduction in TCA intermediates (e.g., malate, citrate) (Fig. 4D). Intriguingly, both N-acetyl-D-galactosamine (GalNAc) and N-acetyl-D-glucosamine (GlcNAc), which are involved in the modification of N-glycosylation and regulate lymphocyte adhesion and maturation, were reduced in DON-treated Tfh cells. Furthermore, b-1,6 N-acetyl-glucosaminyltransferase 5 (*Mgat5*), which initiates GlcNAc- b-1,6 branching at N-glycans, serving as ligands for galectins known to enhance T cell proliferation^37^, was also reduced, suggesting metabolic signals that may control TCR signaling and cell proliferation.

### The inhibition of glutaminolysis in T cells reduced spontaneous autoimmunity

We next investigated the intrinsic role of glutaminolysis in CD4^+^ T cells in lupus with the *Gls1^f/f^* CD4-Cre system (*Gls1* CD4-KO), which has been previously used to investigate glutaminolysis in Th1 and Th17 cells^2^. In addition to the B6 background, we bred *Gls1* CD4-KO to the Sle1b lupus-susceptibility locus, yielding 4 groups of mice *Gls1* CD4-KO and *Gls1^f/f^* (“B6”) controls, and *Sle1b Gls1* CD4-KO and *Sle1b* controls. We first compared the four groups at 9 to 12 months of age, when the *Sle1b* mice show a full expression of autoimmune phenotypes, including the production of anti-dsDNA IgG^31^. *Sle1b Gls1* CD4-KO mice showed a reduced number of splenocytes as compared to *Sle1b* controls, while there was no difference between B6 *Gls1* CD4- KO and B6 mice (Fig. 5A). *Gls1* CD4-KO also reduced the production of anti-dsDNA IgG by *Sle1b* mice (Fig. 5B). *Sle1b* mice present numerous spontaneous large GCs in the spleen that contain numerous GC CD4^+^ T cells, a subset of Tfh cells. *Gls1* CD4-KO reduced the size of *Sle1b* GCs as well as the amount of GC CD4^+^ T cells they contain (Fig. 5C, D).

**Fig. 5.**
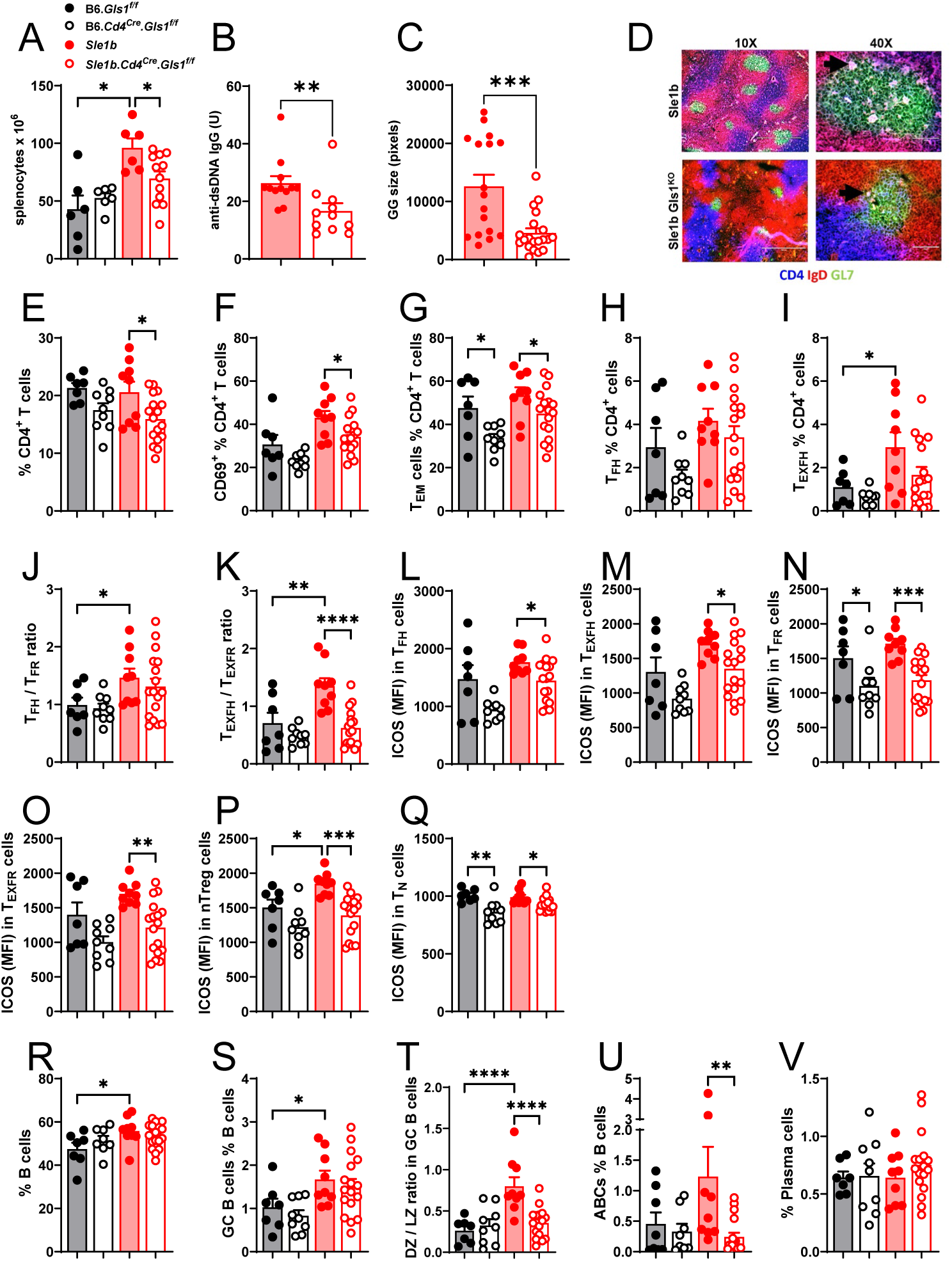
*Gls1* CD4-KO reduced the production of autoantibodies in aged *Sle1b* mice with selective effects on CD4^+^ T cells and B cells. A. Splenocyte number. **B**. Serum anti-dsDNA IgG. **C - D**. Size of GCs in the spleen of 3 mice per group with representative GCs in *Sle1b* (top) and *Sle1b Gls1* CD4-KO (bottom) spleens shown as GL7-positive IgD-negative areas. The 40X images on the right show magenta CD4^+^ T cells inside the GCs. Frequency of CD4^+^ T cells (**E**), CD69^+^ CD4^+^ (**F**), Tem (**G**), Tfh (**H**) Texfh (**I**) cells. Tfh/Tfr (**J**) and Texfh/Texfr (K) ratios. ICOS expression on Tfh (**L**), Texfh (**M**), Tfr (**N**), Texfr (**O**), nTreg (**P**) and Tn (**Q**) cells. Frequency of total B cells (**R**) and GC B cells (**S**). **T**. DZ/LZ ratio in GC B cells. Frequency of ABCs (**U**) and plasma cells (**V**). N = 7 – 17 9 – 12 months old mice. Means ± SEM, compared withMann Whitney tests (B, C) or Šídák’s multiple comparisons tests. *: P < 0.05; **: P < 0.01; ***: P < 0.001; ****: P < 0.0001.

*Sle1b* CD4^+^ T cells showed a higher OXPHOS and glycolysis than B6 CD4^+^ T cells (Fig. S2A, B). *Gls1* CD4-KO did not change these parameters in B6, but it decreased the maximal OXPHOS and glycolysis in *Sle1b* CD4^+^ T cells, suggesting a requirement for glutaminolysis under metabolic stress in these T cells. There was a trend for *Sle1b* CD4^+^ T cells for being more glycolytic than B6, which was accentuated by *Gls1* CD4-KO (Fig. S2C). *Gls1* CD4-KO decreased the frequency of *Sle1b* CD4^+^ T cells as well as their activation evaluated by CD69 expression and Tem cell differentiation (Fig. 5E-G). A similar trend was observed on the B6 background, but it was significant only for the frequency of Tem cells. The frequency of Tfh cells was similar between groups, but *Sle1b* mice presented a higher frequency of Texfh cells with a trend for reduction by *Gls1* CD4-KO (Fig. 5H, I). Both Tfh/Tfr and Texfh/Texfr ratios were higher in *Sle1b* than in B6 mice, and the Texfh/Texfr ratio was reduced by *Gls1* CD4-KO in *Sle1b* mice (Fig. 5J, K), corresponding to the result obtained with DON in W.Yaa mice (Fig. 2H). As in W.Yaa mice treated with DON (Fig. 2I, J and Fig. S1A-D), *Gls1* CD4-KO reduced ICOS expression not only in Tfh-related cells (Fig. 5L-O), but also in Tn and nTreg cells in *Sle1b* mice (Fig. 5P, Q). *Gls1* CD4-KO significantly reduced ICOS expression in B6 mice only in Tfr and Tn cells (Fig. 5N, Q). *Sle1b* mice have higher frequencies of B cells and GC B cells that were not changed by *Gls1* CD4-KO (Fig. 5R, S). Their high DZ/LZ GC B ratio and frequency of ABCs were, however, reduced by *Gls1* CD4-KO (Fig. 5T, V). Finally, contrary to DON-treated W.Yaa mice, there was no effect of *Gls1* CD4-KO on *Sle1b* plasma cells, possibly because their frequency was similar in *Sle1b* and B6 mice (Fig. 5V).

We next evaluated the effect of *Gls1* CD4-KO in young 2 – 3 months old *Sle1b* mice before they produced autoantibodies and in age-matched B6 controls. *Gls1* CD4-KO reduced the frequency of CD4^+^ T cells in both strains (Fig. S3A) to a greater extent than in older mice (Fig. 5E). The frequency of Tfh and Tfr cells was low, as well as the Tfh/Tfr ratio, with no significant difference between strains with and without *Gls1* CD4-KO (Fig. S3B-D). However, the frequency of Texfh and Texfr cells, as well as the Texfh/Texfr ratio, were already higher in *Sle1b* than in B6 mice (Fig. S3E, F), and, as in older mice (Fig. 5K), it was reduced by *Gls1* CD4-KO (Fig. S3G). ICOS expression was also already higher on *Sle1b* Tfh and Texfh cells, and it was downregulated by *Gls1* CD4-KO (Fig. S3H, I). Finally, the higher proliferation of *Sle1b* Tn cells was reduced by *Gls1* CD4-KO (Fig. S3J). *Gls1* CD4-KO increased the frequency of *Sle1b* B cells (Fig. S4K). Young *Sle1b* mice presented a higher frequency of GC B cells in which the high DZ/LZ ratio was reduced by *Gls1* CD4-KO (Fig. S3L, M). The frequency of ABCs and plasmablasts was also higher in young *Sle1b* mice than in B6 mice, with a trend for a reduction of the ABC frequency by *Gls1* CD4-KO (Fig. S3N, O). Finally, there was no difference between groups for the low frequency splenic plasma cells (Fig. S3P).

Overall, these results showed that *Gls1* deficiency in *Sle1b* CD4^+^ T cells phenocopied most of the CD4^+^ T cell phenotypes induced by DON treatment in W.Yaa CD4^+^ T cells, with a most preeminent effect on ICOS expression and the Texfh/Texfr ratio, and with minimal effect on B6 CD4^+^ T cells. These results demonstrate that these phenotypes were targeted by both pharmacological and genetic inhibition of glutaminolysis in a cell-intrinsic manner, in two different lupus-prone genetic backgrounds. *Gls1* deficiency in *Sle1b* CD4^+^ T cells was also sufficient to reproduce the phenotypes obtained in B cells from W.Yaa mice treated with DON, i.e. the reduction of autoantibody production, DZ / LZ GC B cell ratio and the frequency of ABCs. This indicates that these phenotypes are driven in a cell-extrinsic manner by glutaminolysis in T cells.

### The inhibition of glutaminolysis in T cells reduced induced autoimmunity

To further investigate the role of *Gls1* deficiency in T cells in lupus-like autoimmunity, we used the chronic graft vs. host disease model (cGVHD), in which autoantibody production is induced in B6 mice by the transfer of 3-amino acid mismatched H2-b alloreactive CD4^+^ T cells from the B6.bm12 strain^38^. This model largely depends on the extrafollicular activation of autoimmune lymphocytes^39^. One week after induction, *Gls1* CD4-KO mice produced a much-reduced level of anti-dsDNA IgG as compared to B6 mice (Fig. 6A). Although the anti-dsDNA IgG production increased in the subsequent weeks in *Gls1* CD4-KO mice, it was still lower than in B6 mice at week 5 (Fig. 6B), when the spleen immunophenotypes were compared between the two groups. *Gls1* CD4-KO did not have any effect on OXPHOS in the CD4^+^ T cells from these mice, but it increased glycolysis, resulting in a skewed basal OCR / ECAR ratio (Fig. S2D-F). *Gls1* CD4-KO reduced the number of splenocytes and the frequency of Tem cells (Fig. 6C, D). The frequency of IFNγ^+^ CD4^+^ T cells was unchanged, but their number was also decreased by *Gls1* CD4-KO (Fig. 6E, F). As in *Sle1b*, the frequency of Tfh and Texfh cells was not affected with *Gls1* CD4-KO in cGVHD B6 mice (Fig. 6G, H), but ICOS expression was reduced in Texfh cells (Fig. 6I, J). Also, as with *Sle1b* mice, the frequency of GC B cells was unchanged, but there was a trend for a reduced DZ/LZ ratio (Fig. 6K), and more importantly, there was a significant reduction of the frequency of ABCs and plasma cells (Fig. 6L, M). These results showed that *Gls1* CD4-KO induced the same cell-intrinsic effect in CD4^+^ T cells and extrinsic effects in B cells, resulting in a reduction of autoantibody production in the cGVHD induced model of lupus.

**Fig. 6.**
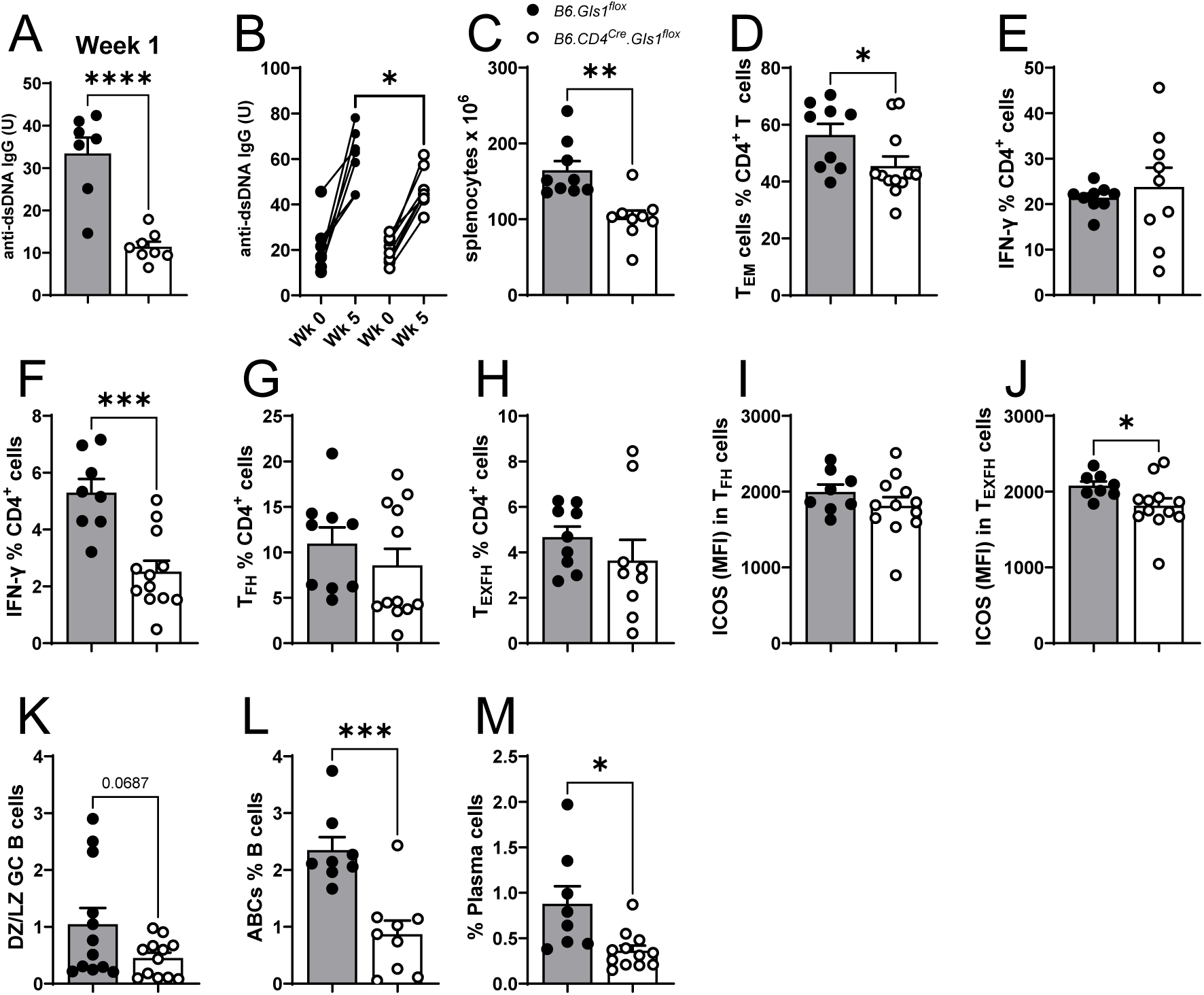
*Gls1* CD4-KO reduced the production of autoantibodies in B6 mice after cGVHD induction with selective effects on CD4^+^ T cells and B cells. A. Serum anti-dsDNA IgG 1 week after induction. **B.** Anti-dsDNA IgG before and 5 weeks after induction. Each pair represents one mouse. **C**-**P**. Phenotypes evaluated 5 weeks after cGVHD induction. **C**. Splenocyte numbers. Frequency of Tem (**D**) and IFNγ^+^ CD4^+^ (**E**) T cells. F. Number of IFNγ^+^ CD4^+^ T cells. Freqeuncy of Tfh (**G**) and Texfh (**H**) cells. ICOS expression on Tfh (**I**) and Texfh (**J**) cells. K. DZ / LZ GC B cell ratio. Frequency of ABCs (**L**) and plasma cells (**M**). N = 9 corresponding to two independent cohorts. Means ± SEM, compared with *t* tests. *: P < 0.05; **: P < 0.01; ***: P < 0.001; ****: P < 0.0001.

### The inhibition of glutaminolysis in T cells altered the humoral response to protein immunization

To further investigate the effect of *Gls1* CD4-KO on T cell help to B cells, we immunized B6 and *Sle1b* mice with and without *Gls1* CD4-KO with a protein antigen, NP-KLH in alum, and boosted them with the same antigen 6 weeks later. Serum high anti-NP4 and low affinity anti-NP25 IgG1, the main isotype elicited by NP immunization, were assessed at weeks 1, 5, and 7. We also measured anti-NP antibodies of the IgG2a isotype, which are elicited by IFNγ-producing CD4^+^ T cells. *Sle1b* mice produced more anti-NP4 and NP-25 IgG1 and IgG2a than B6 mice in the primary response with no difference in affinity, assessed by the NP4/NP25 ratio (Fig. 7A). *Gls1* CD4-KO increased anti-NP IgG1 production in B6 mice leading to a skewed IgG1 to IgG2a ratio (Fig. 6B). In *Sle1b* mice, *Gls1* CD4-KO decreased the affinity of both anti-NP IgG1 and IgG2a at week 5 (Fig. 6C). However, the most striking effect of *Gls1* CD4-KO in *Sle1b* mice was a marked reduction of anti-NP IgG2a, especially at weeks 5 and 7 (Fig. 6C). These results showed that *Gls1* CD4-KO affects the antibody response to immunization with a protein antigen, supporting the hypothesis that the inhibition of glutaminolysis in CD4^+^ T cells alters the help they provide to B cells in a different manner on an autoimmune and healthy genetic background.

**Fig. 7.**
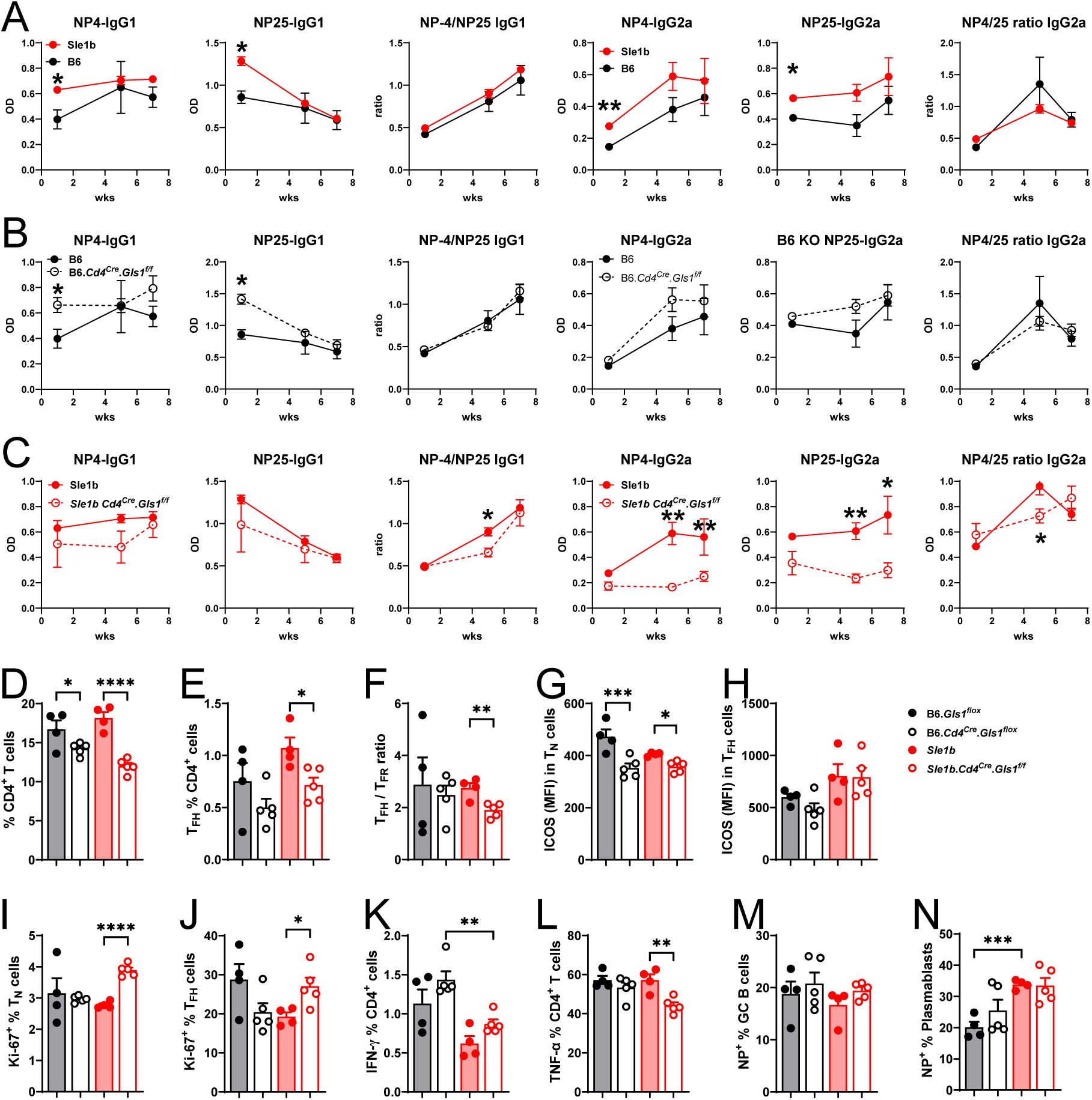
***Gls1* CD4-KO altered antibody response to NP-KLH immunization**. Mice were immunized with NP-KLH and boosted at week 6. **A-C**. Serum anti-NP4 and anti-NP25 IgG1 and IgG2a with NP4/NP25 ratio comparing *Sle1b* and B6 mice (**A**), B6 and B6 *Gls1* CD4-KO (**B**), and *Sle1b* and *Sle1b Gls1* CD4-KO (**C**). Significance is indicated for specific time-points. D–N: Splenocyte analysis at week 7. Frequency of CD4^+^ T cells (**D**) and Tfh cells (**E**)**. F**. Tfh/Tfr ratio. ICOS expression on Tn (**G**) and Tfh (**H**). Frequency of proliferating KI-67^+^ Tn cells (**I**) and Tfh cells (**J**). Frequency of IFNγ^+^ (**K**) and TNFα^+^ (**L**) CD4^+^ T cells. Frequency of NP^+^ in BC B cells (**M**) and plasmablasts (**N**). Means ± SEM, N = 4 – 5 compared with Šídák’s multiple comparisons tests. *: P < 0.05; **: P < 0.01; ***: P < 0.001; ****: P < 0.0001.

Mice were sacrificed at week 7 to evaluate splenic CD4^+^ T cell and B cell phenotypes. As with ex-vivo and cGVHD CD4+ T cells, *Gls1* CD4-KO skewed the basal OCR / ECAR ratio in the CD4+ T cells of immunized mice in favor of glycolysis (Fig. S2G-I).*Gls1* CD4-KO decreased the frequency of total CD4^+^ T cells in both B6 and *Sle1b* backgrounds (Fig. 7D), as well as the frequency of Tfh cells and the Tfh/Tfr ratio in *Sle1b* mice (Fig. 7E, F). ICOS expression was decreased by *Gls1* CD4-KO in Tn cells from both backgrounds, while there was no change in Tfh cells (Fig. 7G, H). This suggested that *Gls1* CD4-KO limited the initiation of the Tfh program in Tn cells. *Gls1* CD4-KO also increased proliferation in *Sle1b* Tn and Tfr cells (Fig. 7I, J). Surprisingly, the frequency of IFNγ^+^CD4^+^ T cells was lower in *Sle1b* than in B6 mice and *Gls1* CD4-KO had no effect (Fig. 7K). This finding contrasts with the higher production of anti-NP IgG2a by *Sle1b* mice and their drastic reduction in *Sle1b Gls1* CD4-KO mice (Fig. 7A-C). The functional kinetics of IFNγ production may have been missed at week 7. The frequency of TNFα ^+^CD4^+^ T cells was reduced by *Gls1* CD4-KO in *Sle1b* mice (Fig. 7L). TNFα production by CD4^+^ T cells increases AID expression in B cells, and class-switching to IgG4, the human equivalent to mouse IgG1^40^. Since *Gls1* CD4-KO had a minimal effect on anti-NP IgG1 production in *Sle1b* mice, the significance of the lower frequency of TNFα^+^ CD4^+^ T cells in *Sle1b Gls1* CD4-KO mice is unclear. At week 7, the frequency of NP^+^ GC B cells was similar between groups (Fig. 7M) and there was no difference in the DZ/LZ ratio (data not shown). *Sle1b* mice showed a higher frequency of NP^+^ plasmablasts, which was not altered by *Gls1* CD4-KO (Fig. 6N). Overall, these results showed that the inhibition of glutaminolysis in T cells altered the antibody response to protein immunization in a different manner between B6 and *Sle1b* mice. Most strikingly, *Gls1* CD4-KO greatly reduced the Th1- driven antigen specific IgG2a production in *Sle1b* mice, while it increased Th2-driven antigen-specific IgG1 in B6 mice.

### The inhibition of glutaminolysis in T cells altered the responses of autoimmune mice to LCMV infection

To specifically determine the effect of *Gls1* CD4-KO on antigen-specific CD4^+^ T cells, we infected mice with the Armstrong strain of the lymphocytic choriomeningitis virus (LCMV) and evaluated the phenotypes of total CD4^+^ T cells as well as LCMV-specific GP66^+^ CD4^+^ T cells in the spleen 7 d later. *Gls1* CD4-KO reduced the frequency of total and GP66^+^ CD4^+^ T cells in *Sle1b* mice (Fig. 8A, B). The same result was obtained for Th1 cells (Fig. 8C, D). Total Tfh cells were also reduced by *Gls1* CD4-KO in *Sle1b* mice, but among Tfh cells, the frequency of GP66^+^ cells was similar between *Sle1b* and *Sle1b Gls1* CD4-KO mice (Fig. 8E, F). *Gls1* CD4-KO also decreased the proliferation of *Sle1b* Th1 cells, but not that of Tfh cells (Fig. 8G, H). We further analyzed GP66^+^ CD4^+^ T cells according to their expression of TCF1, which is intrinsically required for viral-specific TFH cell responses^41^, and CX3CR1, a maker of terminally differentiated Th1 cells in viral infections ^42^. CX3CR1 is also a marker of IFNγ-producing Tfh1 cells^43^, a subset of Tfh cells that is expanded in SLE patients and mouse models of the disease^44–46^. The TCF1^-^ CX3CR1^+^ and, to a greater extent, the TCF1^+^CX3CR1^+^ subsets of GP66^+^ CD4^+^ T cells were decreased by *Gls1* CD4-KO in *Sle1b* mice, while there was no difference for the TCF1^+^ CX3CR1^-^ subset (Fig. 8I-K). The results suggest a greater impact of the inhibition of glutaminolysis on the Tfh1 cells induced by LCMV in *Sle1b* mice. *Gls1* CD4-KO increased the frequency of B cells in *Sle1b* infected mice (Fig. 7L), but among those B cells, it decreased the frequency of GC B cells, as well as the frequency of plasma cells (Fig. 7M, O). Interestingly, LCMV-infected *Sle1b* mice presented a very low frequency of plasmablasts (Fig. 7N), which was the opposite from uninfected mice in the same age-group (Fig. S4O), but it was not affected by *Gls1* CD4-KO. Overall, these results suggest that autoimmune CD4^+^ T cells have a higher requirement for glutaminolysis to respond to LCMV infection as compared to healthy controls. This requirement affects specifically Th1 and Tfh1 cells. Consequently, infected Sle1b *Gls1* CD4-KO mice develop a reduced GC B cell and plasma cell expansion.

**Fig. 8.**
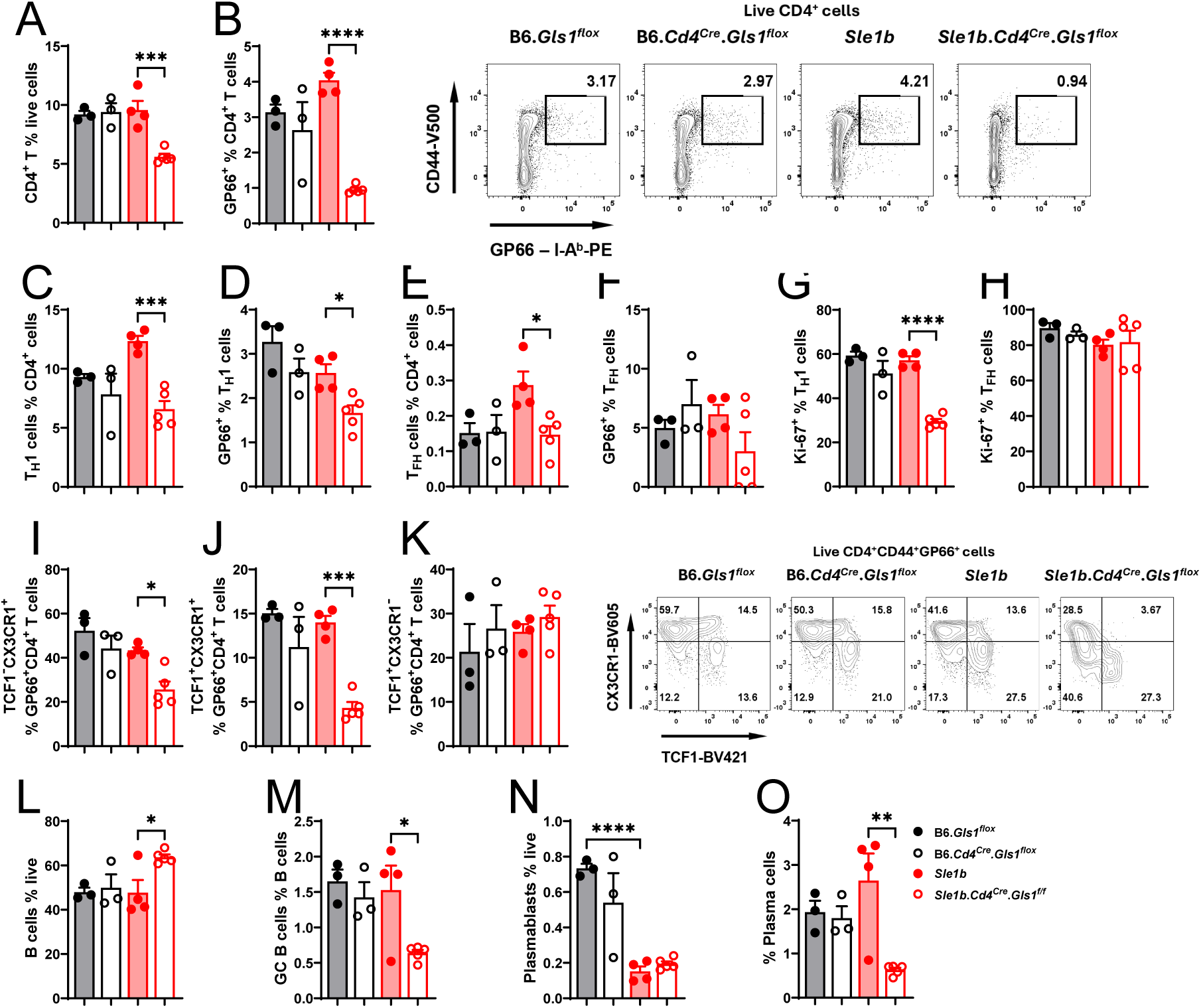
*Gls1* CD4-KO reduced CD4^+^ T cell and B cell response to LCMV infection. Splenic CD4^+^ T cells and B cells were analyzed 7 d after infection with LCMV Armstrong. Frequency of total (**A**) and GP66^+^ (**B**) CD4^+^ T cells, with representative FACS plots on the right. Total (**C**) and GP66^+^ (**D**) Th1 cells, and total (**E**) and GP66^+^ (**F**) Tfh cells. Frequency of KI-67^+^ Th1 (**G**) and Tfh (**H**) cells. Frequency of TCF1^-^CX3CR1^+^ (**I**), TCF1^+^CX3CR1^+^ (**J**), and TCF1^+^CX3CR1^-^ (**K**) GP66^+^ CD4^+^ T cells, with representative FACS plots on the right. Frequency of B cells (**L**), GC B cells (**M**), plasmablasts (**N**) and plasma cells (**O**). Means ± SEM, N = 3 – 5 compared with Šídák’s multiple comparisons tests. *: P < 0.05; **: P < 0.01; ***: P < 0.001; ****: P < 0.0001.

## Discussion

This study focused on the role of glutaminolysis in Tfh cells in lupus-prone mice as compared to healthy controls. Tfh cells are a subset of CD4^+^ T cells that are critical for the production of autoantibodies in SLE^47^, but the role of glutaminolysis has been relatively less studied in Tfh cells comparatively to Th1 and Th17^2, 3, 5^. We showed that the pharmacological inhibition of glutaminolysis with DON reduced the production of anti-dsDNA IgG antibody in TC lupus-prone mice, without changing the frequency of Tfh cells, but reducing their ICOS and BCL-6 expression^16^. The beneficial effect of glutaminolysis inhibition on autoantibody production was confirmed in another lupus-prone MRL/lpr model treated with CB839, along with a modest reduction of Tfh cell frequency^4^. We have shown that glutaminolysis is not required for Tfh cell differentiation from purified CD4^+^ T cells from either lupus or control mice^18^. However, the increased glutamate metabolism and differential expression of glutamine transporters on TC Tfh cells as compared to healthy controls^19^ suggested that glutamine contributes to the expansion of lupus Tfh cells with enhanced functions

Here, we used the TLR7-driven W.Yaa model to further investigate the contribution of glutaminolysis to lupus Tfh cells comparatively to healthy controls. We confirmed that the pharmacological inhibition of GLS1 reduced autoantibody production in this model. The effect was modest, probably due to the treatment lasting 2-weeks only to avoid gastrointestinal toxicity^48^. At the cellular level, we confirmed that GLS1 inhibition reduced the frequency of Th17 cells but not that of Th1^2^, and it had no effect on Treg subsets. DON had also no effect on the frequency of W.Yaa Tfh and Texfh cells. However, DON reduced the expression of ICOS and OX40, which are critical regulators of Tfh cell differentiation, maintenance and function^26,27^. The downregulation of ICOS was observed across all CD4^+^ T cell subsets, including naïve T cells, suggesting that DON may impair their differentiation into Tfh cells. A reduced ICOS expression may also functionally impair Tfh cells. DON-treated and untreated W.Yaa mice presented similar frequency of GC B cells, but DON-treated mice presented a relatively lower number of GC B cells in the dark zone where they migrate after signal received from Tfh cells in the light zone, proliferate and undergo somatic hypermutation before differentiating into plasma cells^49^. The DON treatment also reduced the frequency of plasma cells. The increased TLR7 signaling in W.Yaa mice expands the GC dark zone^50, 51^. In addition, the related *Sle1.Yaa* model shows an altered GC B cell dynamics correlating with plasma cell accumulation and associated with an impaired CD40 signaling in LZ B cells^52^. Our results support the hypothesis that DON either counteracted excessive TLR7 signaling in B cells or altered the function of Tfh cells to restore GC B cell dynamics and their differentiation into plasma cells. DON also reduced the frequency of ABCs, a population greatly expanded in W.Yaa mice. It could be due to a direct effect of DON though the inhibition of glycolysis, since ABCs are highly glycolytic^53^. It could also be due to the reduced Texfh/Texfr ratio and the reduced expression of ICOS on Texfh cells, both functionally reducing their help to ABCs.

DON also uniquely modulated the transcriptome and metabolome of W.Yaa Tfh cells. In addition to the expected downregulation of glutamine metabolism, DON altered the metabolism of other amino acids. DON also reduced the expression of cell cycle genes as genes in the type I IFN pathway, supporting the hypothesis that the TRL7/ type I IFN pathway is sustained by glutaminolysis in W.Yaa Tfh cells. DON had little effect on the transcriptome and metabolome of Tfh cells in B6 mice, which was corroborated by minor phenotypic changes, establishing the unique requirement of lupus Tfh cells for glutaminolysis.

As reported by others^2, 5^, we confirmed that the inhibition of glutaminolysis reduced glycolysis in CD4^+^ T cells, specifically here in lupus Tfh cells, but it has little effect on glycolysis in healthy Tfh cells. The inhibition of glutaminolysis with DON and the inhibition of glycolysis with 2DG^25^ ^22^ had however distinctive effects on W.Yaa Tfh cells, the most striking being that only 2DG decreased their frequency. In addition, only DON reduced N-glycosylation in W.Yaa Tfh cells. 2DG has been reported to reduce N-glycosylation in cancer cells^54^, but there is no evidence that it also occurs in T cells. More detailed metabolic studies will be necessary to dissect the intersection of glycolysis and glutaminolysis in lupus T cells, including N-glycosylation.

To better understand the intrinsic role of glutaminolysis in lupus CD4^+^ T cells, we analyzed the effect of *Gls1*-deficiency in T cells in the autoimmune phenotypes of mice carrying the lupus- susceptibility locus *Sle1b*, which drives the production of autoantibodies associated with an expansion of Tfh cells and GC B cells^28, 29, 30^. We showed that *Gls1* deficiency in T cells was sufficient to reduce autoantibody production, and to phenocopy the effect of DON treatment on Tfh and B cells, including reducing ICOS expression and the Texfh/Texfr ratio, as well as the GC DZ/LZ ratio and the frequency of ABCs. These results were largely confirmed with the cGVHD model of lupus on the B6 background. These results strongly suggest that lupus Tfh cells use glutaminolysis to enhance their help to B cells supporting autoantibody production, in association with altered GC B cell dynamics and an expansion of the ABC subset. We further showed that the inhibition of glutaminolysis in T cells impacted T-dependent humoral responses as well as responses to an acute viral infection in a different manner between an autoimmune and a healthy genetic background. These results support the hypothesis that glutaminolysis supports the function of lupus Tfh cells.

Mechanistically, glutaminolysis may support lupus Tfh cells by promoting the expression of co-stimulatory molecules, especially ICOS. Since ICOS signaling is required for the differentiation and maintenance of Tfh cells by inducing *Bcl6* expression^26^, a higher ICOS expression contributes to Tfh cell expansion. TCR activation triggers *Icos* transcription^55^. W.Yaa Tfh cells express higher levels of *Icos* than B6 Tfh cells, consistent with a stronger TCR signaling^22^. It has been recently shown that *Icos* expression is regulated epigenetically by DNA methylation^56^, a process at least partially regulated by glutaminolysis metabolite αKG. It will be therefore of great interest to assess DNA methylation at the *Icos* promoter in lupus T cells relative to glutaminolysis. Overall, these results have identified a cell-intrinsic role for glutaminolysis in Tfh cells expanded in multiple models on lupus, which indirectly affects B cell differentiation and antibody production. The glutaminase inhibitor CB-839 and a DON-derivative pro-drug are in clinical trials in oncology with promising results^57, 58^. Our study on Tfh cells, combined with other studies on Th17 cells^2, 5^ suggest that these treatments may be beneficial to treat SLE patients.

## Materials and Methods

### Mice and treatment

NZW/J (JAX:001058) females and BXSB/MpJ (JAX:000740) males were bred to produce F1 male progeny (W.Yaa) used in this study. C57BL6/J (B6, JAX:000664) were bred at the UTHSCSA animal facility. Twelve-weeks old W.Yaa and B6 males were treated with intra-peritoneal injections of DON (vendor, 1.6 mg /kg) or solvent controls 3 times a week for 2 weeks.

B6.*Gsl1f/f* mice were bred to CD4-Cre mice (JAX:017336) as well as B6.*Sle1b* mice^31^ to produce 4 strains B6.*Gsl1f/f* (B6), B6.*Gsl1f/f-CD4-Cre* (*Gls1* CD4-KO), B6.*Sle1b* (*Sle1b*), and B6.*Sle1b.Gsl1f/f-CD4-Cre* (Sle1b *Gls1* CD4-KO). Only females were used from these strains at either 2-3 month or 9-12 months of age.

Chronic graft-versus-host disease (cGVHD) was induced as previously described^38^. Briefly, B6.*Gsl1f/f* and B6.*Gsl1f/f-CD4-Cre* hosts received 8 x 10^7^ splenocytes from B6.H-2bm12 mice (JAX #001162) via intra-peritoneal injection. Sera were collected weekly after induction and stored for ELISA measurement of autoantibodies. Hosts were sacrificed 5 weeks after transfer and splenocytes were analyzed by flow cytometry.

For dinitrophenyl-keyhole limpet hemocyanin (NP-KLH) immunization, 8–10- week-old mice received an intraperitoneal injection of 100 μg NP(31)-KLH (Biosearch Technology) in alum as previously described^16^, and were boosted with the same immunogen at week 6. Mice were sacrificed at week 7 to evaluate splenic CD4^+^ T cell and B cell phenotypes. Some mice in the same age group were infected intraperitoneally with 0.5 x 10^6^ PFU LCMV-Armstrong and sacrificed 7 d later for immunophenotyping of splenocytes.

Mice were maintained in SPF conditions at the University of Texas Health San Antonio (UTHSA). This study was carried out in accordance with the guidelines from the Guide for the Care and Use of Laboratory Animals of the Animal Welfare Act and the <u>NIH (National Research Council) Guide</u> <u>for the Care and Use of Laboratory Animals</u>. All animal protocols were approved by the Institutional Animal Care and Use Committee of UTHSA (IACUC 20220032AR).

### Antibody measurement

Anti-dsDNA IgG were detected by ELISA in sera diluted 1:100 as previously described^16^. Antinuclear antibodies (ANAs) were detected on Hep-2 cell-coated slides in sera diluted 1:40. NP- specific antibodies were measured in the serum of NP-KLH immunized mice collected at weeks 1, 5 and 7 after the primary immunization. ELISA plates were coated with NP(4)- or NP(25)-BSA (high or low affinity, respectively) (Biosearch Technology), followed by incubation with 1:5,000 (IgG1) or 1:1,000 (IgG2a) diluted serum samples, and developed with alkaline phosphatase-conjugated goat anti-mouse IgG1 (1071-04, 1:1000) or IgG2a (1081-04, 1:1000), both from Southern Biotech. All samples were run in duplicate.

### Flow cytometry and cell sorting

Flow cytometry was performed on splenocyte suspensions prepared as previously described^16^. Dead cells were excluded with fixable viability dye. For intracellular staining, cells were fixed and permeabilized using the FOXP3/Transcription Factor Staining Buffer. Data were acquired using an LSRFortessa or FACSymphony A5, and analyzed with FlowJo V10 (BD Bioscience). Gating strategies for CD4^+^ T cell and B cell subsets are shown in Figure S4A. Tfh cells were gated as FOXP3^-^CD44^+^PD-1^+^CXCR5^+^BCL6^+^PSGL1^lo^CD4^+^ T cells, Tfr cells as FOXP3^+^PD- 1^+^CXCR5^+^BCL6^+^PSGL1^lo^CD4^+^ T cells, Texfh cells as FOXP3^-^CD44^+^PD-1^+^CXCR5^-^BCL6^-^PSGL1^lo^CD4^+^ T cells and Texfr cells as FOXP3^+^CD44^+^PD-1^+^CXCR5^-^BCL6^-^ PSGL1^lo^CD4^+^ T cells.

In the LCMV infection experiment, Th1 cells were gated as CD44^+^PD-1^-^PSGL1^+^CD4^+^ T cells. I-A(b) LCMV GP 66-77 DIYKGVYQFKSV (GP66) tetramers were obtained from the NIH Tetramer core facility. For RNASeq and metabolomic analyses, Tfh cells were sorted as CD4^+^CD44^+^PD- 1^+^PSGL-1^lo^ cells with a FACSARIA III cytometer. Purity was > 95% (Figure S4B).

### RNA-seq

Tfh cells were FACS-sorted from W.Yaa and B6 mice treated with DON or untreated controls. RNA was isolated from purified cells using a RNeasy Plus Micro kit (Qiagen) and cDNA libraries were constructed using a SMART-seq HT (Takara), as previously described ^59, 60^. Paired-end sequence reads (2 x 150 bp) were aligned to the mouse reference genomes (GRCm38) using STAR v2.7.5c. Normalized counts, represented by transcripts per million (TPM), were generated using RSEM v1.3.3. The raw transcript counts were used as input to determine significantly expressed genes by DESeq2 based on the criteria (TPM > 1, FDR < 0.05, fold change >1.5). Gene set enrichment analysis (GSEA) was performed using DAVID (https://david.ncifcrf.gov/).

### Metabolomic analysis

Mitochondrial stress assays were conducted on an XF96 extracellular flux analyzer on magnetic-bead purified CD44^+^ CD4^+^ T cells or CD4^+^ T cells, as indicated for each experiment. Data was normalized using a coupled Biotek Cytation 1 imager. Purified Tfh cells (same samples used for RNA-seq analysis) were processed for metabolomic analysis as described previously ^59, 61^. Metabolic feature alignment and curation were performed by MZmine. After normalizing to total ion chromatogram, intensities were tested for group significance using unpaired Student *t*-test. Metabolites were identified by comparison to the metabolomic library of purified standards. Metabolic pathway analysis was performed using *Mummichog.* The pathways represented by at least 3 significant metabolites are presented.

### Statistical analysis

Statistical analyses were performed using GraphPad Prism 9.0 software. Unless otherwise stated, differences between groups were evaluated by ANOVA with correction for multiple testing or unpaired or paired *t* tests, as indicated in the figure legends. The corresponding nonparametric tests were used when the data distribution deviated from normality. Results were expressed as means ± standard errors of the mean (SEM). Statistical significance levels were set at *: *P* < 0.05, **: *P* < 0.01, ***: *P* < 0.001 and ****: *P* < 0.0001.

| KEY RESOURCES TABLE |  |  |  |  |  |
| --- | --- | --- | --- | --- | --- |
| Antibodies and stains |  |  |  |  |  |
| Target Antigen | Conjugation | Clone | vendor | Dilution | RRID # |
| BCL-6 | BV750 | K112-91 | BD Biosciences | 1:50 | AB_3146163 |
| CD4 | BUV395 | GK1.5 | BD Biosciences | 1:100 | AB_2738426 |
|  | BV750 |  | BioLegend | 1:100 | AB_2734150 |
|  | AF700 |  | Thermo Fisher | 1:100 | AB_494001 |
| CD5 | AF700 | 53-7.3 | BioLegend | 1:100 | AB_2687002 |
| CD11c | APC | N418 | Thermo Fisher | 1:100 | AB_469346 |
| CD16/CD32 |  | 2.4G2 | BD Biosciences | 1:500 | AB_394656 |
| CD19 | BV786 | 1D3 | BD Biosciences | 1:100 | AB_2738141 |
| CD21 | BV711 | 7E9 | BioLegend | 1:100 | AB_2876440 |
| CD23 | BV510 | B3B4 | BioLegend | 1:100 | AB_2563705 |
| CD25 | BV421 | PC61 | BD Biosciences | 1:100 | AB_11153485 |
| CD44 | PE | IM7 | BioLegend | 1:100 | AB_312959 |
|  | V500 |  | BD Biosciences | 1:100 | AB_312959 |
| CD62L | APC | MEL-14 | BD Biosciences | 1:100 | AB_398533 |
|  | AF700 |  | Thermo Fisher | 1:100 | AB_494003 |
| CD69 | PE-Cy7 | H1.2F3 | BioLegend | 1:100 | AB_493564 |
| CD86 | BV650 | GL-1 | BioLegend | 1:100 | AB_11126147 |
| CD93 | BV750 | AA4.1 | BD Biosciences | 1:100 | AB_2871654 |
| CD95 (Fas) | Biotin | Jo2 | BD Biosciences | 1:100 | AB_395328 |
| CD134 (OX-40) | BV650 | OX-86 | BD Biosciences | 1:100 | AB_2740249 |
| CD138 | BV605 | 281-2 | BioLegend | 1:100 | AB_2562336 |
| CD162 (PSGL-1) | BV605 | 2PH1 | BD Biosciences | 1:100 | AB_2740114 |
|  | AF647 |  | BD Biosciences | 1:100 | AB_2737807 |
| CD184 (CXCR4) | BUV395 | 2B11 | BD Biosciences | 1:100 | AB_2740007 |
| CD185 (CXCR5) |  | 2G8 | BD Biosciences | 1:100 | AB_394302 |
| CD278 (ICOS) | PE | 7E.17G9 | BD Biosciences | 1:100 | AB_394349 |
| CD279 (PD-1) | eFluor 450 | RMP1-30 | Thermo Fisher | 1:100 | AB_11150068 |
|  | BV785 | J4329F.1A12 | BioLegend | 1:100 | AB_2563680 |
| CX3CR1 | BV605 | SAO11F11 | BioLegend | 1:100 | AB_2565999 |
| FOXP3 | FITC | FJK-16S | Thermo Fisher | 1:100 | AB_465243 |
| GL7 | eFluor450 | GL7 | Thermo Fisher | 1:100 | AB_10870775 |
| GP66 tetramer | APC |  | NIH | 1:15 |  |
| Helios | AF700 | 22F6 | Thermo Fisher | 1:100 | AB_2784773 |
| IgD | FITC | 11-26c.2a | BD Biosciences | 1:100 | AB_394859 |
| IL-10 | PE-Cy7 | JES5-16E3 | BioLegend | 1:100 | AB_11150582 |
| IL-17A | PE | TC11-18h10.1 | BioLegend | 1:100 | AB_315464 |
| IFN-γ | BV421 | XMG1.2 | BioLegend | 1:100 | AB_893526 |
| Ki-67 | BV711 | SolA15 | BD Biosciences | 1:100 | AB_2738406 |
| NP | PE | B56 | Biosearch Tech | 1:100 | AB_2738406 |
| T-bet | PE-Cy7 | eBio4B10 | Thermo Fisher | 1:100 | AB_2784726 |
| TCF1 | BV421 | S33-966 | BD Biosciences | 1:100 | AB_2869822 |
| Streptavidin | PerCP |  | BD Biosciences | 1:100 |  |

## ACKNOWLEDGMENTS

We thank the UF SECIM metabolomic core for technical support, the staff of the UTHSA Flow Cytometry Shared Resource, as well as Dr. Nu Zhang (UTHSCSA) for assistance with LCMV experiments.

## AUTHOR CONTRIBUTIONS

Conceptualization, S.C.C., M.M., and L.M. Methodology, S.C.C. and Y.G., Investigation, S.C.C, Y.G., M.V.J., D.J., L.T.P., and C.L.; Writing – Review & Editing, M.M. and L.M.; Resources, J.C.R.; Funding Acquisition, M.M., and L.M.; Supervision, L.M.

