## supplemental table and figures for "Glutaminolysis promotes the function of follicular helper T cells in lupus-prone mice"

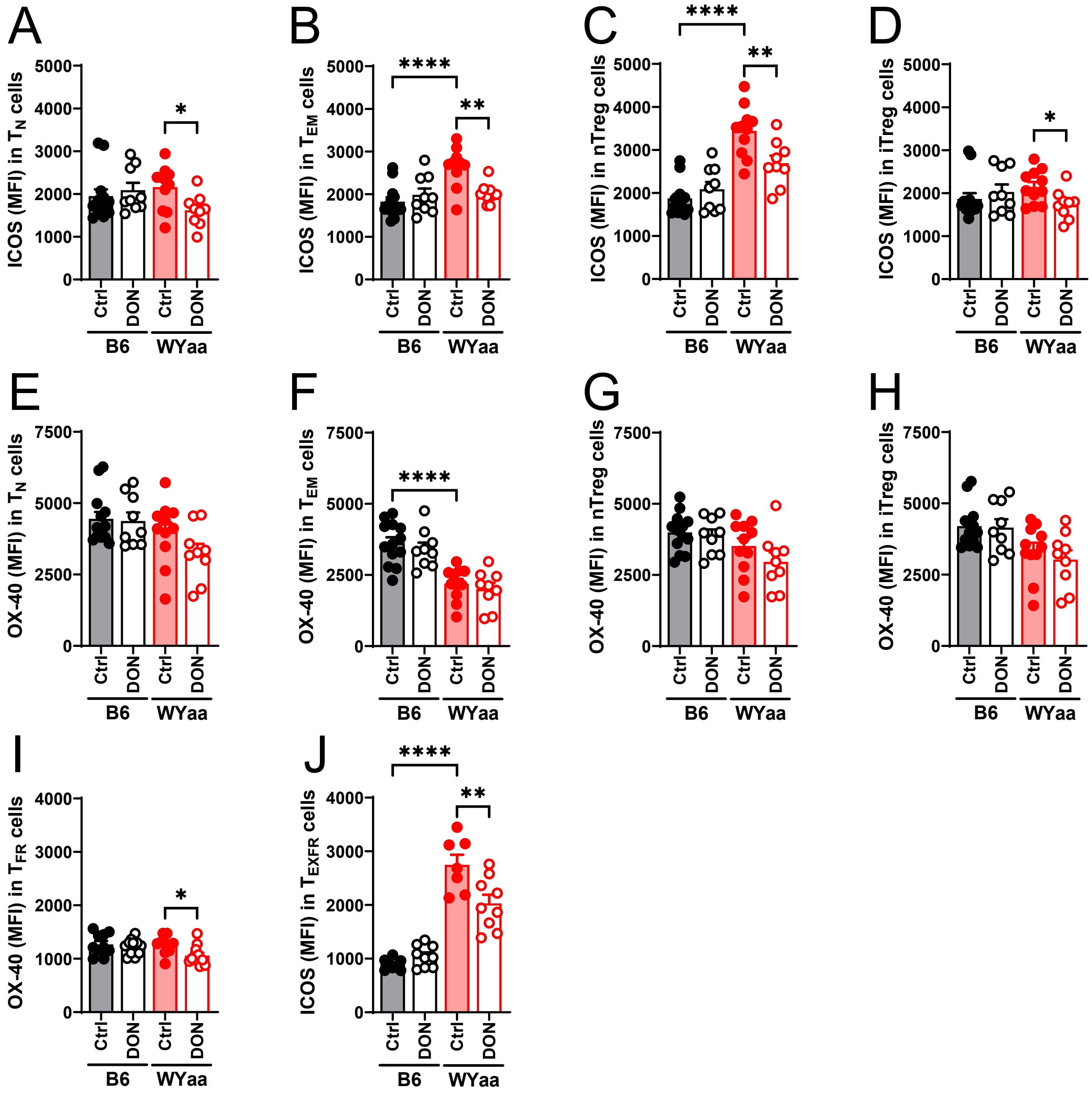


**Fig. S1. DON reduced ICOS and OX-40 expression.** ICOS expression on Tn (**A**), Tem (**B**), nTreg (**C**) and iTreg (**D**) cells. OX-40 expression on Tn (**E**), Tem (**F**), nTreg (**G**), iTreg (**H**) cells, Tfr (**I**) and Texfr (**J**) cells. Means $\pm$ SEM, N = 10 – 15 compared with Šídák's multiple comparisons tests. *: P < 0.05; **: P < 0.01; ***: P < 0.001; ****: P < 0.0001.

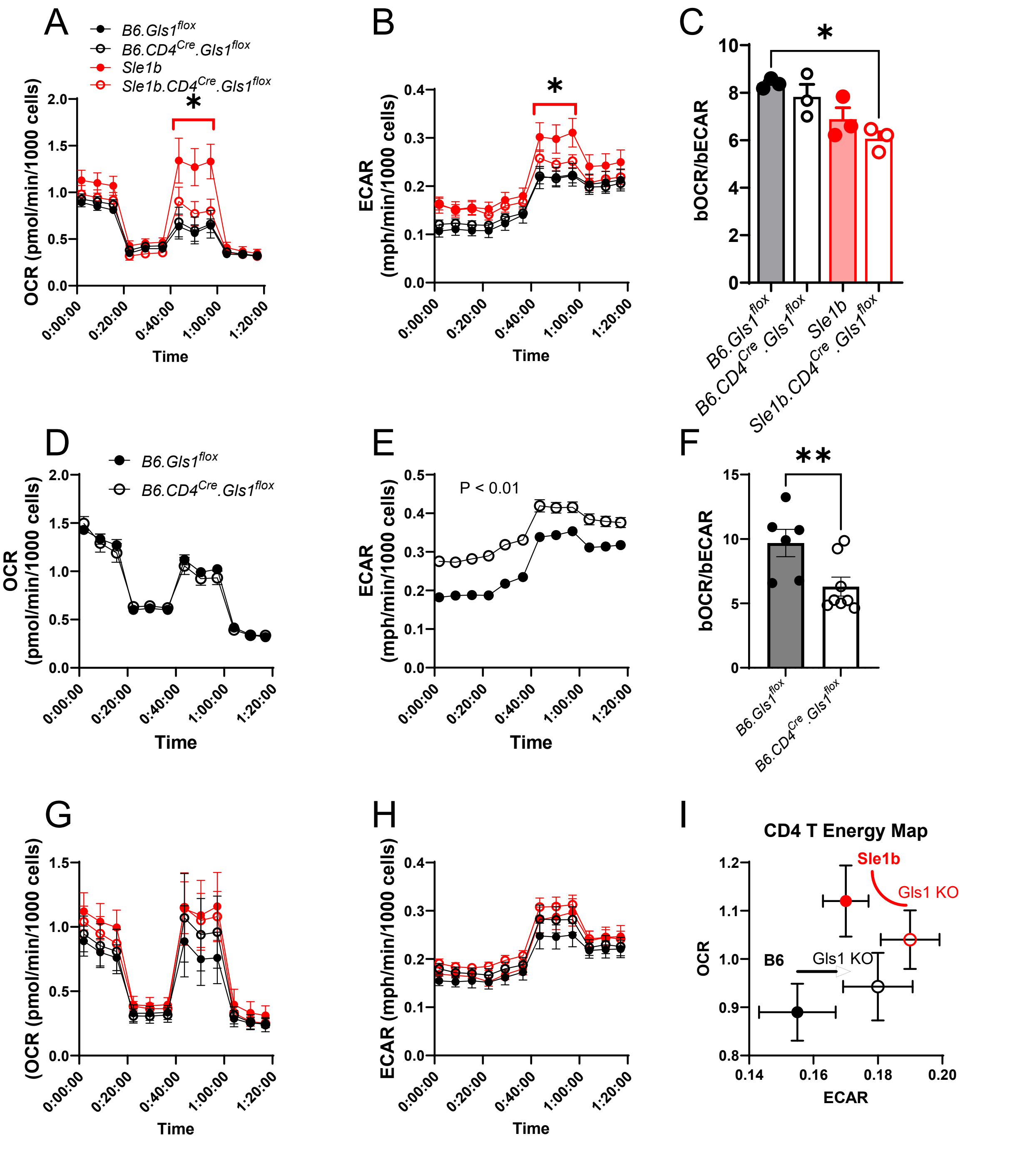


**Fig. S2. *Gls1* CD4-KO increased glycolysis in CD4⁺ T cells.** Mitochondrial stress assays were performed on CD4⁺ T cells from untouched 9-12 months old mice (**A-C;** N = 3), 5 weeks after cGVHD induction (**D-E,** N = 6 - 8), and 7 weeks into a TD-recall response to NP-KLH immunization (**G-I,** N = 5). OCR (**A**, **D**, **G**) and ECAR (**B**, **E**, **H**). bOCR/bECAR ratio (**C**, **F**). **I**. Energy map. Mean + SEM. **A** and **B**: t tests for the 3 time points in the brackets between Sle1b and *Sle1b Gls1* CD4-Cre. **E**. t tests for each time point. **C** and **F**: Mann-Whitney tests.

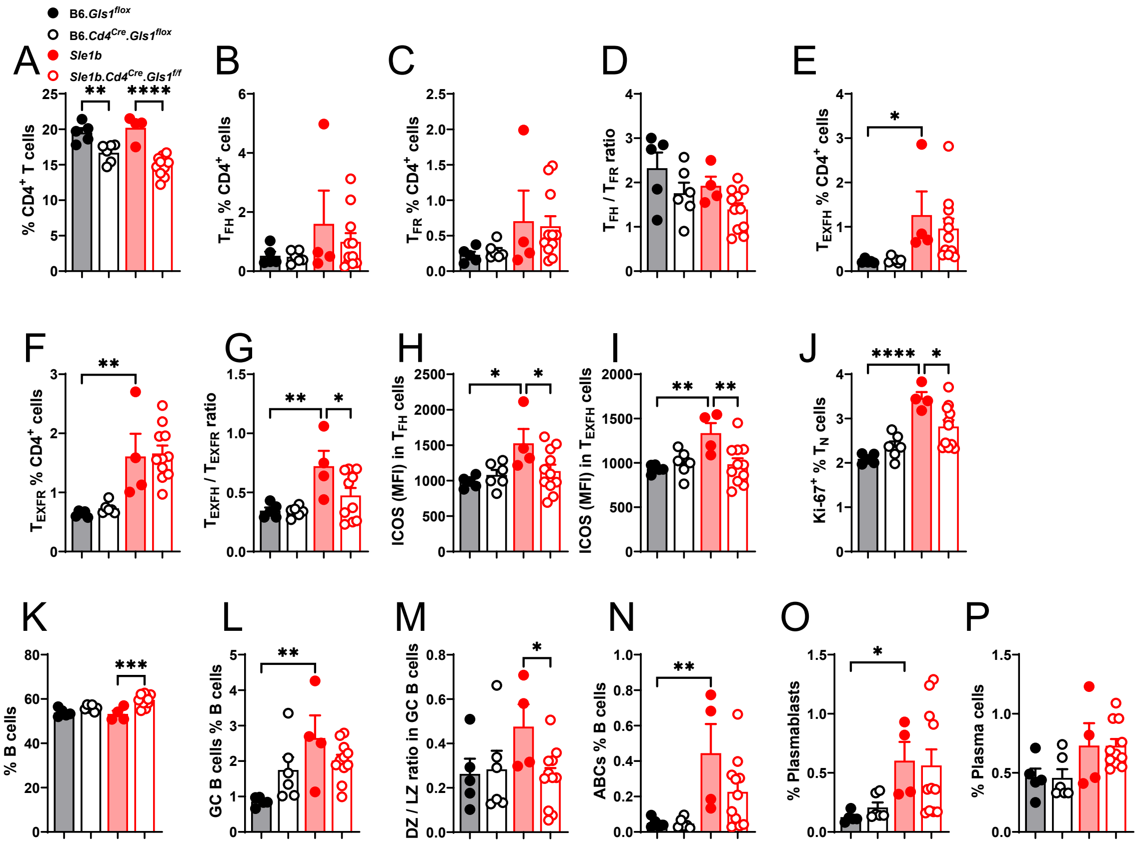


**Fig. S3. The alteration of immunophenotypes of *Sle1b* mice by *Gls1* CD4-KO were initiated before autoantibody production.** Frequency of CD4$^{+}$ T cells (**A**), Tfh (**B**), Tfr (**C**) cells and Tfh/Tfr ratio (**D**). Frequency of Texfh (**E**), Texfr (**F**) cells and Texfh/Texfr (**G**) ratio. ICOS expression on Tfh (**H**) and Texfh (**I**) cells. (**J**) Frequency of Ki-67$^{+}$ Tn cells. Frequency of B cells (**K**) and GC B cells (**L**). **M**. DZ/LZ ratio in GC B cells. Frequency of ABCs (**N**), plasmablasts (**O**) and plasma cells (**P**). N = 4 – 11 2-3 months old mice. Means $\pm$ SEM, compared with Šídák's multiple comparisons tests. *: P < 0.05; **: P < 0.01; ***: P < 0.001; ****: P < 0.0001.

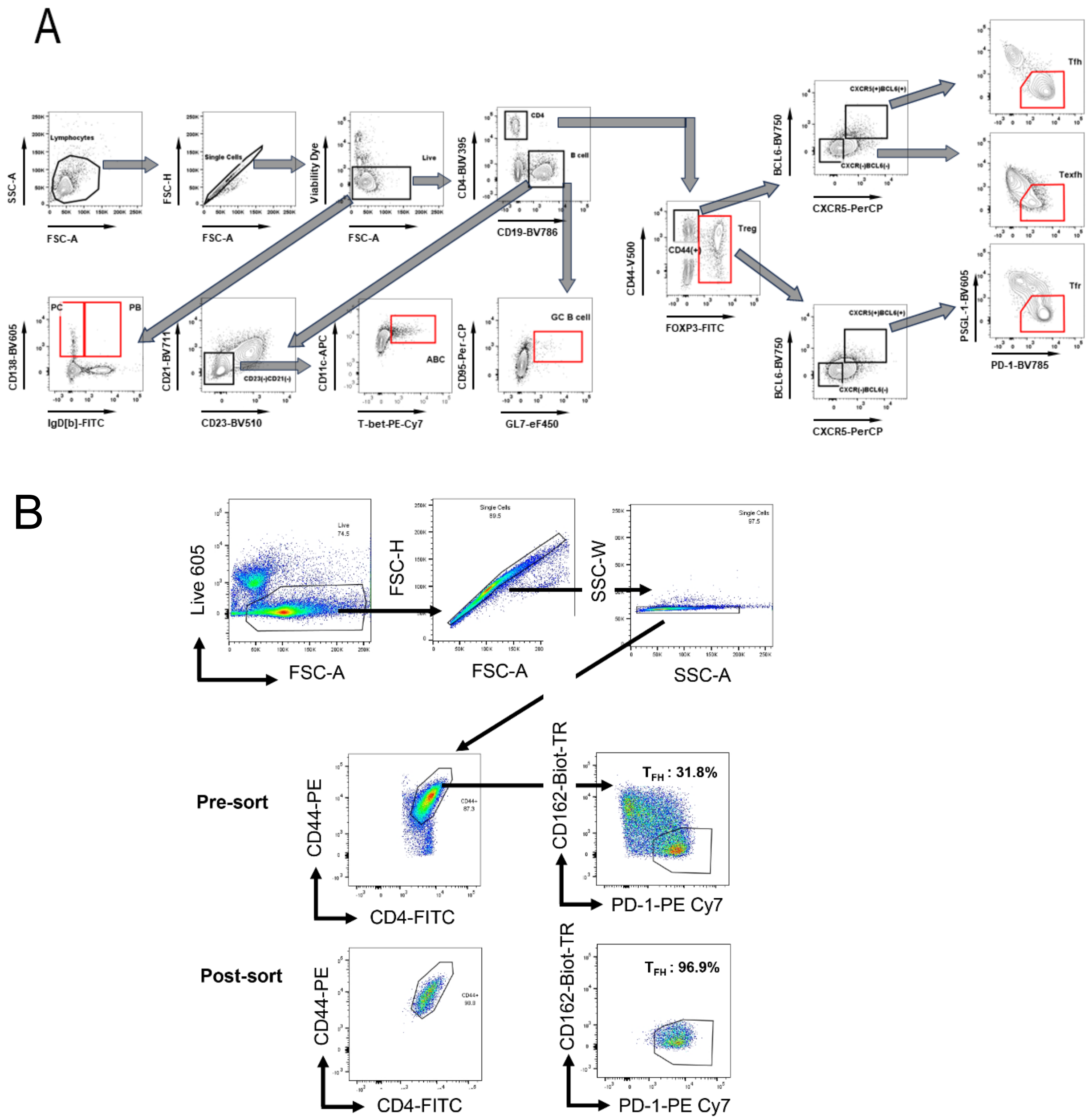


**Fig.S4**. **A**. Gating strategies for mouse CD4$^+$ T cells and B cells. **B**. Representative FACS plots showing sorting strategy and efficiency of $T_{FH}$ cells. Purified splenic CD4$^+$ T cells were sorted into $T_{FH}$ cells as CD44$^+$ PSGL-1 (CD162)$^{lo}$ PD-1$^+$. The pre- and post-sort gates are shown for both cell types.
